# Blind but alive - congenital loss of *atoh7* disrupts the visual system of adult zebrafish

**DOI:** 10.1101/2024.04.23.590799

**Authors:** Juliane Hammer, Paul Röppenack, Sarah Yousuf, Anja Machate, Marika Fischer, Stefan Hans, Michael Brand

**Affiliations:** CRTD - Center for Regenerative Therapies at TU Dresden, Dresden, Germany

## Abstract

1

**Purpose:** Vision is the predominant sense in most animal species. Loss of vision can be caused by a multitude of factors resulting in anatomical as well as behavioral changes. In mice and zebrafish, *atoh7* mutants are completely blind as they fail to generate retinal ganglion cells during development. In contrast to mice, raising blind zebrafish to adulthood is challenging and this important model is currently missing in the field. Here, we report the phenotype of homozygous mutant adult zebrafish *atoh7* mutants that have been raised using adjusted feeding and holding conditions.

**Methods:** The phenotype of adult mutants was characterized using classical histology and immunohistochemistry as well as optical coherence tomography. In addition, the optokinetic response was characterized.

**Results:** Adult *atoh7* mutants display dark body pigmentation and significantly reduced body length. They fail to form retinal ganglion cells, the resulting nerve fiber layer as well as the optic nerve, and consequently behave completely blindly. In contrast, increased amounts of other retinal neurons and Müller glia are formed. In addition, the optic tectum is anatomically reduced in size, presumably due to the missing retinal input.

**Conlusions:** Taken together, we provide a comprehensive characterization of a completely blind adult zebrafish mutant with focus on retinal and tectal morphology, as a useful model for glaucoma and optic nerve aplasia.

## 2 Introduction

Vision is the predominant sense in most vertebrate species including humans ^1,2^. It is primarily dependent on the eye’s structures, notably the retina. The lens focuses incoming light onto the retina, a stratified neural tissue ^3^. Within the retina’s outer nuclear layer (ONL), photoreceptor cells capture this light and transform it into signals. These signals are subsequently transmitted to bipolar cells in the inner nuclear layer (INL). Interconnected amacrine and horizontal cells further modify these signals and relay them to retinal ganglion cells (RGCs) in the ganglion cell layer (GCL) which then serve as projection neurons to retinofugal targets in the brain via the optic nerve (ON).

However, this complex system is susceptible to disturbances, causing vision impairment. Globally, millions of people are affected by conditions ranging from minor visual impairments to complete blindness ^4^. Particularly, hereditary disorders that impede RGC and ON development can lead to irreversible vision loss ^5^. Beside rodents, zebrafish have emerged as invaluable models for studying these disorders ^6–9^. Like all vertebrates, zebrafish have a highly conserved retinal structure, and they possess 4 different types of cone photoreceptors (red, green, blue and UV) ^10,11^. Similar to humans, zebrafish are diurnal and their retina is dominated by cones. They rely heavily on vision, e.g. for prey detection, predator avoidance and shoaling behavior. Their visual functionality can be assessed as early as four days post-fertilization using optomotor (OMR) or optokinetic response (OKR) tests ^12,13^. In these assays, larvae are shown a moving stimulus, such as a grating pattern. A functioning visual system is indicated if larvae follow the stimulus direction either by swimming (OMR) or by eye movement (OKR) ^7,12,14^.

A large number of visual mutants have been identified by alterations in these visually-mediated behaviors ^7,14^,^15,16^; specifially, the *lakritz* mutant *(lak^th241^)* was identified due to its dark pigmentation in the large Tübingen screen for zebrafish mutants ^15,17^. Normally, zebrafish are able to adapt their body pigmentation to ambient light levels mediated by melanophores; the *lakritz* mutant, however, displays a permanently dark body pigmentation, reminiscent of licorice (*Lakritz* is German for licorice). Additionally, it was shown that larvae produce no positive reaction in OMR and OKR tests, indicating blindness ^14,18^. Morphological analyses revealed a notably thinner GCL, with only about 20% of cells remaining, while the INL was slightly enlarged ^19^. The cause for these morphological defects was found to be a single nucleotide mutation in the atonal-homolog 7 locus (*atoh7*, formerly atoh5) ^19^, a vertebrate homologue of atonal, originally identified for its crucial role in chordotonal sense organ development in *Drosophila* ^20^. In healthy retinae, development occurs in sequentially timed, but overlapping waves: the proliferative neuroepithelium initially produces RGCs, followed by cells of the INL, and finally PRs and Müller glia ^21^. Studies in larval zebrafish have shown that mutations in the *atoh7* gene results in a failure of retinal progenitor cells to develop into RGCs, and therefore absence of the optic nerve. Instead of RGCs, an excess amount of INL neurons are produced ^19^. *Atoh7*, a bHLH transcription factor central to a regulatory gene network affecting retina development, is closely linked with various pathways and is highly conserved across vertebrates, including humans ^22,23^. Mutations in *atoh7* in humans have been associated with multiple ocular diseases, including optic nerve hypoplasia and aplasia, disorders of the retinal vasculature and glaucoma ^24–26^.

Although a large number of larval and also some adult visual mutants have been extensively described and characterized (e.g. ^7,9^), data on fully blind zebrafish models are scarce, especially in adult stages. To our knowledge, the only documented adult blind zebrafish mutant is bumper (bum) ^27^. Bumper mutants are visually impaired resulting from malformation and ectopic placement of the lens. Although their blindness is well characterized in larvae, characterization in adults relies on qualitative morphologic and behavioral assessments. The complete failure of RGC development in the *atoh7* mutant results in a unique neuronal environment with the brain being completely disconnected from the visual system. Recent studies have analyzed the impact of the lack of retina-derived signals on the larval brain ^28^, however, it would be of great interest to also study these effects in adults. Here, we introduce a paradigm for raising *atoh7* mutants to adulthood, allowing comprehensive characterization of the adult phenotype. We find that the larval reduction of RGCs can not be compensated for in homozygous adults, resulting in lack of the ON and complete blindness in adult *atoh7* zebrafish, also at the behavioral level. Additionally, we identified morphological changes in visual processing areas of the midbrain that are the likely indirect consequence of retinofugal input. We suggest that the adult *atoh7* mutant described here provides a valuable tool to study vision, visual processing and regenerative approaches for the visual system, providing a new experimental model for glaucoma and optic nerve aplasia in zebrafish.

## 3 Results

### 3.1 *Atoh7* mutants can be raised to adulthood and show a distinct phenotype

In order to obtain homozygous *atoh7* mutants, heterozygous *atoh7^+/-^*fish were crossed and progeny were raised to adulthood. However, under standard raising conditions ^29^, we did not obtain any adult homozygous mutant fish. Hence, the raising strategy was modified and mutant fish were raised separately to enhance their survival (Fig. 1A). To select for homozygous mutants, the offspring was screened for a dark pigmentation (caused by the impaired background adaptation) at 7 days post fertilization (dpf). While the majority of larvae displayed a normal pigmentation, about 25 % of the larvae were characterized by the typical darker pigmentation pattern (Fig. S1A). To confirm the genotype, individual homozygous larvae were genotyped by PCR (Fig. S1B and Methods). Sequencing of the PCR products confirmed the genotyping strategy (Fig. S1D). Sorted larvae were raised to adulthood with light and dark phenotype being raised separately but under comparable conditions. In brief, fish were raised in smaller groups with an excess supply of food and delayed connection to the aquatic system (see also Methods). The overall survival of the mutants was above 50 % (53.5 %), but still lower compared to siblings (81.5 %) (Fig. 1B). Adult mutants displayed a characteristic phenotype as they retained their characteristic dark pigmentation to adulthood (Fig. 1C). In addition, mutant fish showed a decelerated growth during development that resulted in a significantly reduced body length at 6 month post fertilization (mpf) (*atoh7^+/+^:* 35.8 ± 0.9 mm*, atoh7^+/-^:* 35 ± 1 mm*, atoh7^-/-^:* 28.7 ± 1.3 mm) (Fig. 1D). Mutant fish were sexually mature and can produce viable offspring, albeit apparently at a reduced fertilization rate (not shown). During all experiments, heterozygous *atoh7* fish displayed a comparable phenotype to the wildtype siblings and are therefore not specifically mentioned during the subsequent descriptions.

**Figure 1:**
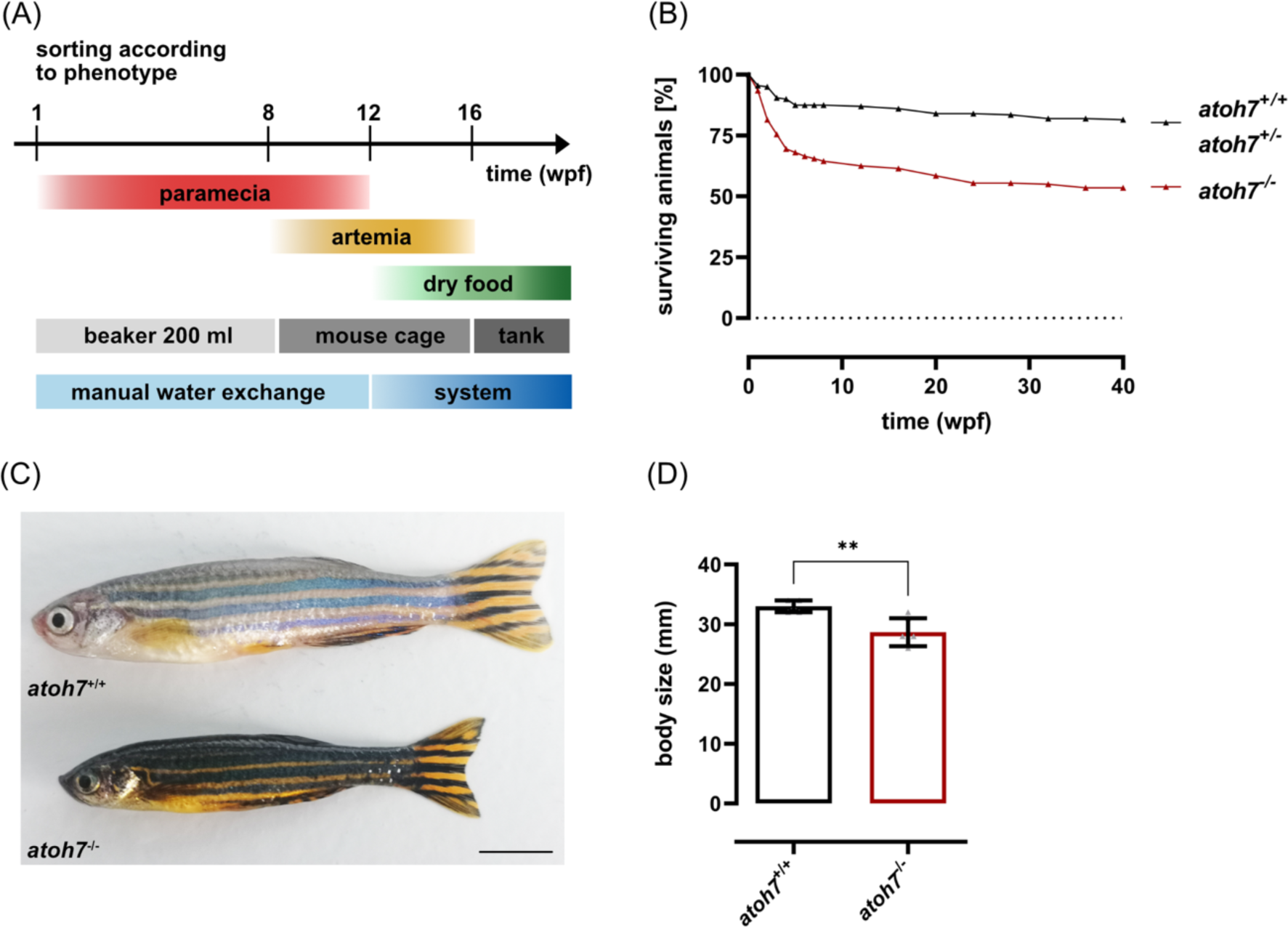
Adult *atoh7* mutants display a dark body pigmentation and growth retardation. (A) Scheme of the modified raising strategy to obtain adult mutant animals. Mutants and wildtype/heterozygous siblings are separated according to their phenotype at 1 wpf (Fig. S1) and raised under the same conditions. Until 8 wpf, animals are kept in 200 ml beakers with manual water exchange every other day and exclusively fed with paramecia. At 8 wpf, animals are transferred into mouse cages and additionally fed with artemia. At 12 wpf, mouse cages are connected to the system. Feeding of paramecia is terminated and more and more complemented with dry food, in addition to paramecia. By 16 wpf, animals are exclusively fed with dry food and moved to tanks. (B) Quantification of the survival of wildtype/heterozygous versus mutant fish revealed a lower survival rate of mutant animals. (C) Adult mutants (6 months post fertilization) are characterized by a darker pigmentation as well as a reduced body length. (D) Quantification of the body size revealed a significantly reduced length of mutants compared to wildtype and heterozygous siblings. Scale bars represent 5 mm. Statistics: all data are represented as mean ± SD, one-way ANOVA with Tukey’s multiple comparison test, p < 0.05 (*); 0.01 (**); 0.001 (***) or 0.0001 (****). Abbreviations: wpf – weeks post fertilization.

### 3.2 Adult *atoh7* mutants are blind due to the lack of retinal ganglion cells

To obtain insights into the structure and morphology of the retina of mutant fish, we performed hematoxylin-eosin (HE) stainings on retinal cross sections of wildtype and mutant retinae. Initial examination revealed that mutants have an overall intact eye structure including lens and layered retina (Fig. 2A). Retinal layering was assessed in more detail with the help of HE stainings as well as optical coherence tomography (OCT) *in vivo* recordings ^30,68^. The wildtype retina showed the characteristic layering with all three nuclear layers. Although the three nuclear layers and the corresponding plexiform layers were also present in the mutant, the retinal nerve fiber layer (RNFL) could not be detected. In addition, the ganglion cell layer (GCL) appeared less densely populated and photoreceptor outer segments seemed less organized. As the mutant retina appeared thinner in the histological sections and in the *in vivo* OCT recordings, the thickness of the total retina as well as of the individual layers was measured. The basis for these measurements were *in vivo* OCT images, as the fixation and processing steps in HE sections might cause tissue shrinkage. The absolute thickness of the mutant retina was significantly reduced compared to wildtype (*atoh7^-/-^:* 193. ± 8.5 µm, *atoh7^+/+^:* 271.2 ± 8 µm, n = 6)) (Fig. S2A) as well as the thickness of all layers except the outer plexiform layer (Table S3, Fig. S2B). As mutants were smaller, we normalized the retinal thickness in relation to body length, thus allowing a more accurate comparison between mutants and wildtype siblings. Although the relative retinal thickness was still significantly reduced in the mutant, the difference was not as pronounced as in absolute values (*atoh7^+/+^:* 8.2 ± 0.3 RU*, atoh7^-/-^:* 6.8 ± 0.7 RU, n ≥ 5, Fig. 2B). For the individual layers, thickness was normalized to the total retinal thickness, and was not significantly different (Table 1, Fig. 2C). Hence, the mutant retina is significantly thinner due to the absent retinal nerve fiber layer (RNFL). Next, we quantified the number of cells in the GCL, and found a significant reduction in mutants (*atoh7^+/+^:* 23.9 ± 4.1 µm*, atoh7^-/-^:* 15.3 ± 2 µm, n = 6) (Fig. 3A). To determine the identity of these cells, we analyzed the expression of *islet2b*, a known marker of mature RGCs ^30^, using qPCR, and found a highly significant downregulation in mutant retinae (*atoh7^+/+^:* 1 ± 0.026, *atoh7^-/-^:* 0.14 ± 0.018, n = 3) (Fig. 3B). To confirm the lack of RGCs in mutant fish further, we performed an *in situ* hybridization against *islet2b* (Fig. 3C). While wildtype retinae showed *islet2b* expression in the GCL (indicated by accumulation of yellow dots) only background staining was observed in the mutants. We therefore conclude that the remaining cells in the mutant GCL are not RGCs.

**Figure 2:**
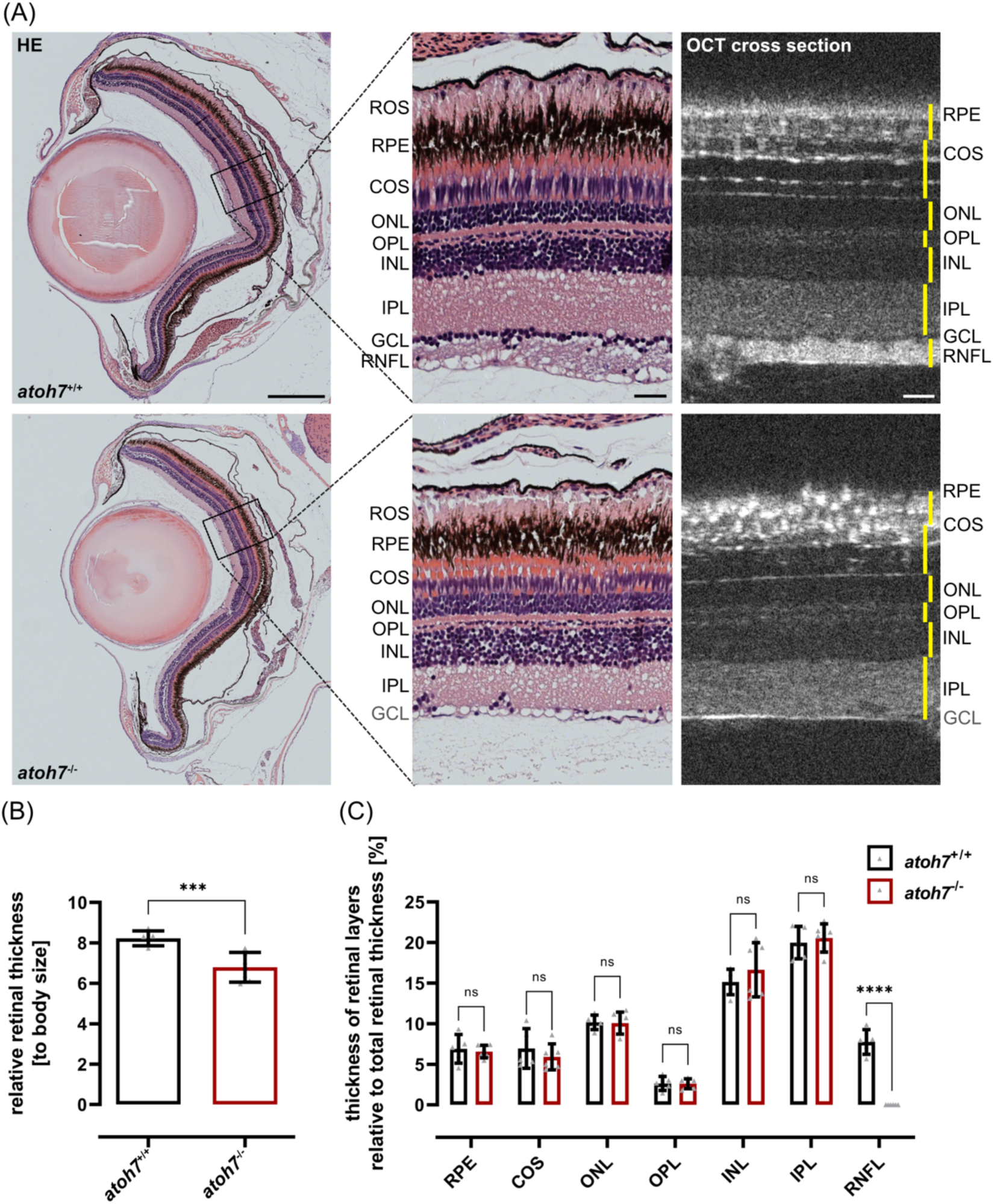
The retinal thickness is reduced in *atoh7* mutants due to a missing nerve fiber layer. (A) HE stainings of wildtype and mutant animals. Overall, retinal morphology is similar in wildtype as well as mutants and individual nuclear and plexiform layers can be distinguished (left panel). However, detailed analysis (middle panel) revealed that mutant retinae miss the retinal nerve fiber layer (RNFL). Additionally, the cone outer segments (COS) appear to be less organized and the rod outer segments (ROS) are shorter in mutant specimens. Optical coherence tomography (OCT) cross sections in live animals confirmed these observations (right panel). (B) Quantification showed that the relative retinal thickness of mutant fish (compared to body size) is significantly reduced. (E) Quantifications of the relative thicknesses of individual layers (as % of whole retinal thickness, measured layers depicted in yellow in (A)) showed no significant differences except for the RNFL. Scale bars represent 250 µm in overviews and 25 µm in magnifications. Statistics: all data are represented as mean ± SD, unpaired t test (B, D), t test with multiple comparisons (C, E), p < 0.05 (*); 0.01 (**); 0.001 (***) or 0.0001 (****). Abbreviations: COS – cone outer segments, GCL – ganglion cell layer; INL – inner nuclear layer, IPL – inner plexiform layer; ONL – outer nuclear layer, OPL – outer plexiform layer; RNFL – retinal nerve fiber layer; ROS – rod outer segments, RPE – retinal pigment epithelium.

**Figure 3:**
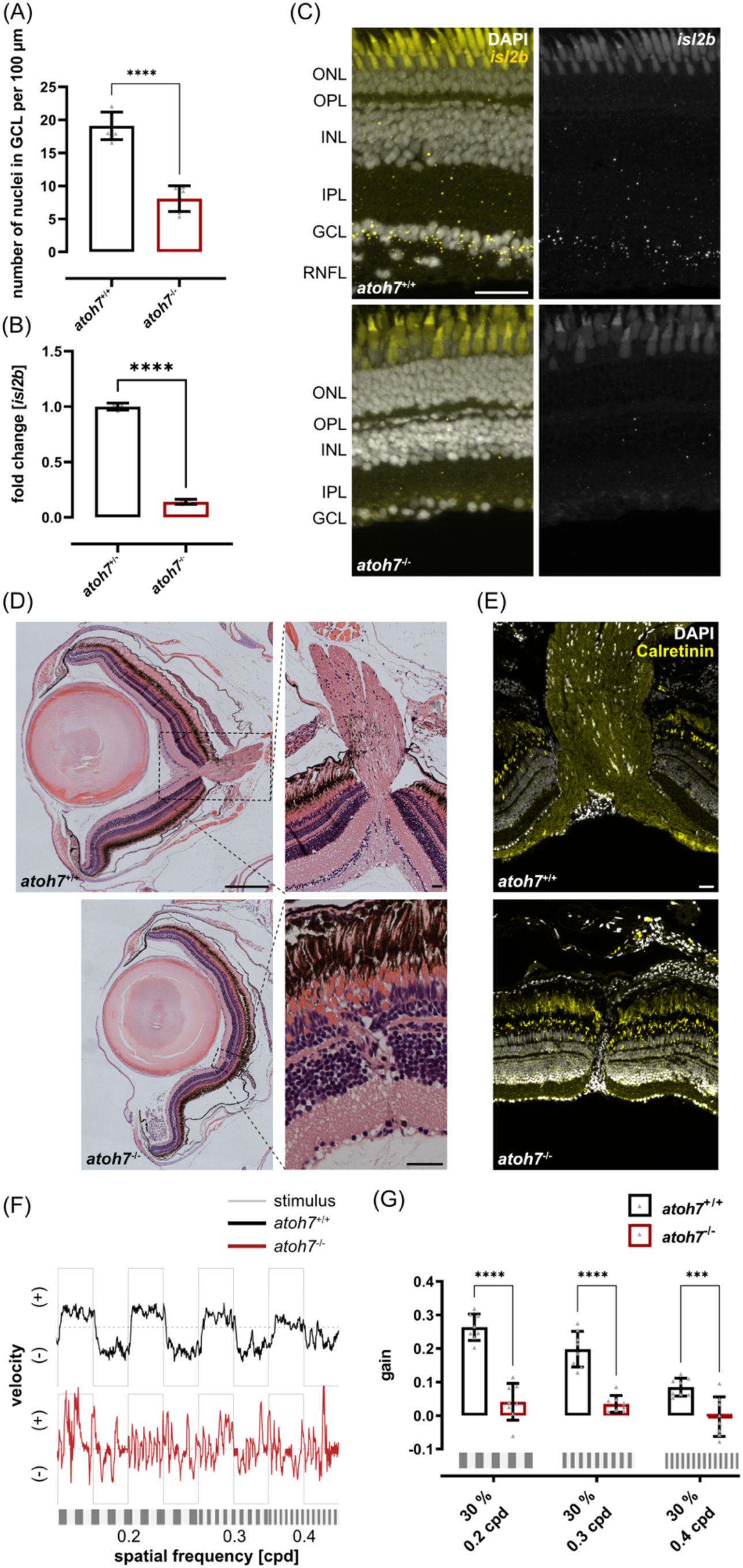
*Atoh7* mutants lack retinal ganglion cells and are blind. (A) Quantification of the number of nuclei in the GCL revealed a significant reduction in mutant fish. (B) qPCR analysis showed a highly significant reduction of the ganglion cell specific marker islet2b mRNA in mutant retinae. (C) Expression analysis of islet2b using in situ hybridization shows specific expression (yellow dots) in the GCL of wildtype but not in mutant retinae. Nuclei were stained with DAPI (white). (D) Representative cross sections of wildtype and mutant retinae stained with HE at the level of the optic nerve. In contrast to wildtype fish that form a regular optic nerve (upper panel), the optic nerve is absent in mutant retinae which rather contain an ectopic blood vessel connecting the subretinal space with the vitreous body (lower panel). (E) Calretinin (yellow), a marker for nerve fibers, is only detected in wildtype but not in mutant retinae. Nuclei were stained with DAPI (white). (F) Graphic representation of the velocity in anti-clockwise (+) and clockwise (-) direction of the eye movements of a representative wildtype (black line) and mutant (red line) fish depending on the stimulus (grey line). Wildtype fish track the stimulus reliably at low spatial frequencies in both directions but movements become more random at high spatial frequencies. Only random eye movements could be observed in mutant fish. (G) The optokinetic response (OKR) was measured with varying spatial frequencies (black-white, 30 % contrast, 0.2/0.3/0.4 cycles per degree (cpd)). While *atoh7*+/+ and *atoh7*+/- achieved comparable gains dependent on the spatial frequency (values), no optokinetic response could be observed in mutants. Scale bars represent 250 µm in overviews and 25 µm in close-ups. Statistics: all data are represented as mean ± SD, t test with multiple comparisons, p < 0.05 (*); 0.01 (**); 0.001 (***) or 0.0001 (****). Abbreviations: cpd – cycles per degree. Abbreviations: GCL – ganglion cell layer; INL – inner nuclear layer; IPL – inner plexiform layer; ONL – outer nuclear layer; OPL – outer plexiform layer; RNFL – retinal nerve fiber layer; RPE – retinal pigment epithelium.

**Table 1:** Absolute and relative thicknesses of retinal layers in *atoh7* mutants and wildtypes.

| layer | absolute thickness [μm] |  |
| --- | --- | --- |
|  | *atoh7*<sup>+/+</sup> | *atoh7*<sup>-/-</sup> |
| RNFL | 21.1 ± 4.1 | na |
| IPL | 54.2 ± 4.9 | 40.7 ± 2.6 |
| INL | 41.1 ± 4.2 | 33.6 ± 5.6 |
| OPL | 7.1 ± 2 | 4.8 ± 1 |
| ONL | 27.5 ± 1.7 | 20.5 ± 1.6 |
| POS | 18.9 ± 6.1 | 12.2 ± 2.7 |
| RPE | 18.7 ± 4.1 | 2.5 ± 1 |

All samples n = 6. Abbreviations: INL – inner nuclear layer; IPL – inner plexiform layer; ONL – outer nuclear layer; na – not available; OPL – outer plexiform layer; POS – photoreceptor outer segments, RNFL – retinal nerve fiber layer, RPE – retinal pigment epithelium.

As anticipated from this finding, no optic nerve was present in histological sections. Instead, we observed a single blood vessel entering the retina from the subretinal space and branching into the vitreal space in all analyzed mutant retinae (Fig. 3D). We used Calretinin, a marker for nerve fibers, to further investigate the makeup of the RNFL. In wildtype specimens, antibody stainings showed the presence of Calretinin-positive nerve fibers throughout the RNFL and the optic nerve (Fig. 3E). Conversely, in the mutant, Calretinin staining was absent, indicating the lack of nerve fibers and thus, of retinotectal projections.

To characterize visual function of the adult *atoh7* mutants, we measured the optokinetic response (OKR) (Fig. 3F,G) ^31^. The fish were presented with black-white stimuli with a contrast of 30 % and decreasing spatial frequencies (0.2 to 0.4 cpd). While wildtype fish were able to track the stimuli, as can be inferred from the correlation between directed eye movements into the direction of the stimulus, we observed only erratic eye movements in the mutants. Independent of testing conditions, the gain (ratio of the tracking speed of the eyes to the speed of the stimulus) observed in mutants was very low, indicating that no OKR was elicited and hence mutants can be considered blind (Table 2). In contrast, the measured gain in wildtypes was significantly higher and dependent on the presented stimuli.

**Table 2:** Individual data points for the OKR test in *atoh7* mutants.

| SF<br>[cpd] | gain |  |  |
| --- | --- | --- | --- |
|  | *atoh7*<sup>+/+</sup> | *atoh7*<sup>+/-</sup> | *atoh7*<sup>-/-</sup> |
| 0.2 | 0.26 ± 0.04 | 0.25 ± 0.04 | 0.04 ± 0.05 |
| 0.3 | 0.02 ± 0.05 | 0.22 ± 0.04 | 0.03 ± 0.02 |
| 0.4 | 0.08 ± 0.02 | 0.1 ± 0.03 | 0 ± 0.06 |

All samples n = 8. cycles per degree – cpd, SF – spatial frequency. Abbreviations: AV – angular velocity; cpd – cycles per degree; dps – degrees per second; SF – spatial frequency. AV 15 dps, contrast 30 %.

### 3.3 Excess numbers of inner retinal cells are produced in the mutant retina

To examine the identity of the remaining cells in the GCL, we performed antibody stainings against HuC/D, a specific marker for RGCs and most amacrine cells (Fig. 4A). In both wildtype and mutant retinae, HuC/D-positive cells were detected in both GCL and INL. Although not quantified, the vast majority of cells in the GCL of both samples were HuC/D-positive. The quantification of the total number of HuC/D-positive cells per 100 µm retina revealed no significant difference between wildtype and mutants (*atoh7^+/+^:* 25 ± 3, *atoh7^-/-^:* 24 ± 2, n = 6) (Fig. 4B). However, in mutants, there were significantly fewer HuC/D-positive cells in the GCL (*atoh7^+/+^:* 10 ± 2, *atoh7^-/-^:* 5 ± 1, n = 6) (Fig. 4C) and significantly more in the INL (*atoh7^+/+^:* 15 ± 2, *atoh7^-/-^:* 19 ± 2, n = 6) (Fig. 4D). Due to expression of HuC/D but lack of *islet2b*, we assumed that the cells in the GCL of the mutants are of amacrine identity. Next, to explore the effect of missing *atoh7* expression on other retinal cell types, we used immunohistochemical markers. To analyze bipolar cells, we used the marker PKCα. In wildtype retinae, the cell bodies of PKCα-positive cells resided within the INL, extending processes to the OPL and GCL (Fig. 4E). Although the morphology of bipolar cells remained unchanged between wildtype and mutant specimens, a significant increase in their numbers was observed in mutants. (*atoh7^+/+^:* 13 ± 1, *atoh7^-/-^:* 19 ± 2 per 100 µm retina, n = 6) (Fig. 4G). Next, we analyzed Müller glia using GFAP (glial acidic fibrillary protein) as a marker. In wildtype retinae, the cell bodies of GFAP-positive Müller glia could be detected in the INL and their processes spanned the entire retina from GCL to the photoreceptor layer (Fig. 4F). The morphology of Müller glia again was unaltered in mutants, but their number was significantly increased compared to wildtypes (*atoh7^+/+^:* 12 ± 1, *atoh7^-/-^:* 15 ± 1, n = 6) (Fig. 4H).

**Figure 4:**
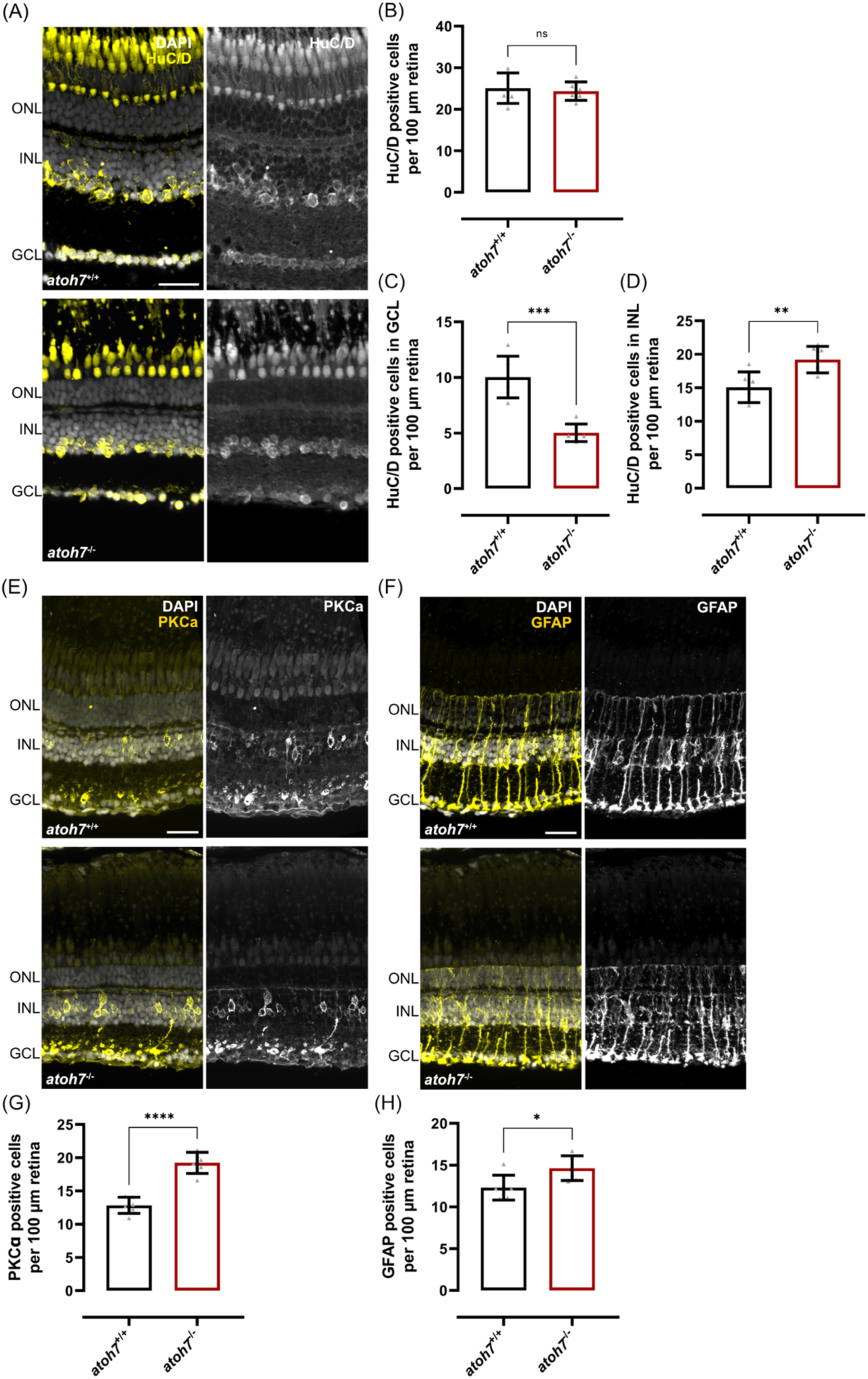
Mutant retinae contain excess numbers of amacrine and bipolar cells as well as Müller glia. (A) Representative images of antibody stainings against HuC/D (yellow), a marker for amacrine and ganglion cells, show HuC/D expression in the INL and GCL of both mutant and wildtype retinae. Nuclei are stained with DAPI (white). (B) Quantification of the overall number of HuC/D positive cells in both INL and GCL revealed no statistical difference. (C,D) Quantifications of HuC/D positive cells in individual layers indicated a significant reduction in the GCL (C) and a significant increase in the INL of mutant retinae (D). (E) Representative images of bipolar cells labeled with PKCα (yellow) showed that cell bodies are located in the INL and GCL of both mutant and wildtype retinae. Nuclei were stained with DAPI (white). (G) Quantification of PKCα positive cells showed a significant increase in mutant retinae. (F) Representative images of an antibody staining against the Müller glia cell specific marker GFAP (yellow) shows cell bodies of Müller glia cells that were located in the inner nuclear layer (INL) of wildtype and mutants retinae as well as their processes that span the entire retina. Nuclei were stained with DAPI (white). (H) Quantification of zrf1-positive cells revealed a significant increase in mutant retinae. Scale bars represent 25 µm. Statistics: all data are represented as mean ± SD, unpaired t test, p < 0.05 (*); 0.01 (**); 0.001 (***) or 0.0001 (****). Abbreviations: GCL – ganglion cell layer; INL – inner nuclear layer; ONL – outer nuclear layer.

### 3.4 The structure of the midbrain shows alterations in *atoh7* mutants

As shown above, adult mutants do not possess an optic nerve. Consequently, the mutant brain never receives any retinal input. To determine the effect of missing retinal input on the brain, we analyzed the structure of the whole brain and, more specifically, the optic tectum – as the main retinorecipient area^32^. All observable brain areas were present in the mutant as depicted in dorsal and ventral top views of representative animals (Fig. 5A). In direct comparison, the mutant brain appeared reduced in size, however, the normalized brain area showed no significant difference (*atoh7^+/+^:* 571 ± 24, *atoh7^-/-^:* 571 ± 38, n = 6) (Fig. S3A). To detect potential differences in the size of individual brain areas, we calculated the relative size of telencephalon, optic tectum and cerebellum (dashed outlines) in comparison to the sum of the three areas (Fig. 5B). While the area of the optic tectum was reduced in size in the mutant (*atoh7^+/+^:* 46.7 ± 2.4 %, *atoh7^-/-^:* 19 ± 2, n = 6), the area of the telencephalon was significantly larger (*atoh7^+/+^:* 27.7 ± 5.1 %, *atoh7^-/-^:* 35.8 ± 2.4 %, n = 6). The size of the cerebellum, however, was similar between wildtype and mutants (*atoh7^+/+^:* 22.5 ± 3.1 %, *atoh7^-/-^:* 23.8 ± 2 %, n = 6). Next, we investigated possible structural changes in the mutant optic tectum via representative coronal brain sections stained with HE (Fig. 5C). To ensure accurate comparisons, anatomical landmarks were used to select sections from similar regions. The overall structure of the optic tectum was preserved in the mutants, however, individual parts appeared to be slightly changed in size. To quantify these differences, the absolute areas of optic tectum, periventricular grey zone and tectal ventricle were normalized to the area of the brain stem including the tegmentum (here designated as Teg) which was not changed in size in the mutants (Fig. S3B,C). The relative area of the optic tectum was significantly reduced in mutants (*atoh7^+/+^:* 0.86 ± 0.12 RU, *atoh7^-/-^:* 0.61 ± 0.1 RU, n = 4). In contrast, the relative size of the periventricular grey zone was not changed, despite appearing thicker in the HE stained sections (*atoh7^+/+^:* 0.27 ± 0.04 RU, *atoh7^-/-^:* 0.23 ± 0.03 RU, n ≥ 4) (Fig. 5D,E). The area of the tectal ventricle was also highly significantly reduced in the mutants (*atoh7^+/+^:* 0.15 ± 0.05, *atoh7^-/-^:* 0.03 ± 0.02, n ≥ 4) (Fig. 5F). Also of note were significant alterations in the layering of the mutant optic tectum. Typically, the optic tectum is comprised of six distinct layers, as clearly identifiable in histological sections from wildtype retinae (Fig. 5G). In the mutants, the stratum opticum could not be detected and the stratum fibrosum et griseum superficiale was significantly thinner in mutants (*atoh7^+/+^:* 63.46 ± 3.78 RU, *atoh7^-/-^:* 54.69 ± 4.68 RU, n = 8) (Fig. 5H).

**Figure 5:**
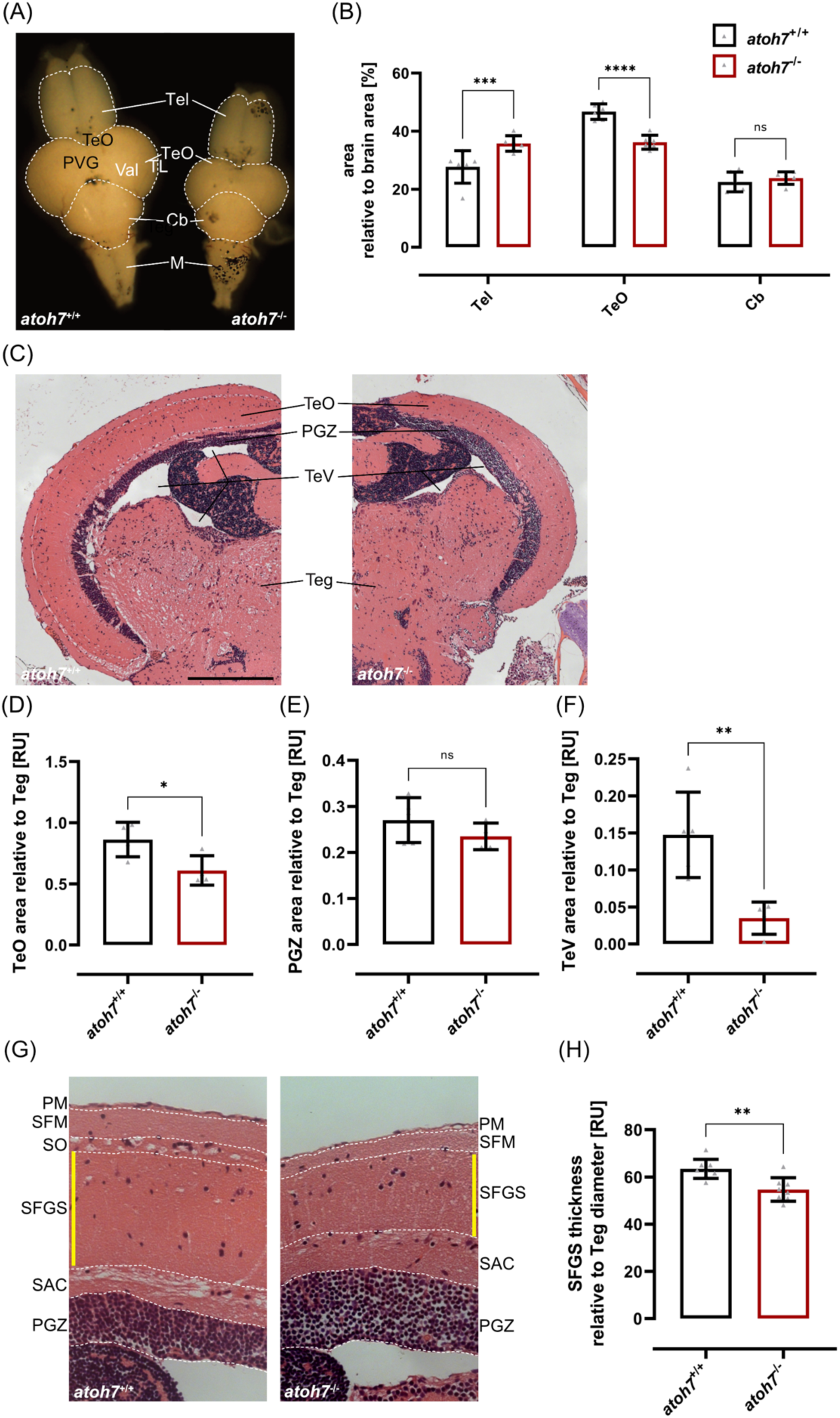
Midbrains of *atoh7* mutants are characterized by structural changes in the area of the optic tectum. (A) Dorsal views of representative wildtype and mutant brains. (B) Comparison of the relative area of specific brain parts compared to the area of the whole brain reveals that mutants possess a significantly larger telencephalon (Tel) but a significantly smaller optic tectum (TeO). In contrast, the cerebellum (Cb) displays no difference. (C) Representative cross sections of the optic tectum of wildtype and mutant stained with HE. The TeO appears reduced in size and the PGZ appeared thicker. The tectal ventricle (TeV) was also smaller in the mutant brain. (D) Quantifications confirmed that the area of the TeO (relative to the area of the Tegmentum (Teg)) was significantly reduced in mutants. (E) The relative area of the PGZ (normalized to Teg area) was not altered in mutants compared to wildtypes. (F) Quantification of the relative area of the TeV (normalized to the Teg area) indicated a very strong reduction in mutants. (G) Representative cross sections of the optic tectum stained with HE. In wildtype, six layers can be distinguished: pia mater (PM), stratum fibrosum marginale (SFM), stratum opticum (SO), stratum fibrosum et griseum superficiale (SFGS), stratum album centrale (SAC) and stratum periventriculare (SPV). In the mutant, no stratum opticum (SO) could be detected and stratum fibrosum et griseum superficiale (SFGS) appeared to be thinner. (H) Quantification of the relative thickness of the SFGS (normalized to Teg diameter) confirmed a significant reduction in mutants. Scale bars represent 250 µm in overviews and 25 µm in close-ups. Statistics: all data are represented as mean ± SD, unpaired t test, p < 0.05 (*); 0.01 (**); 0.001 (***) or 0.0001 (****).

## 4 Discussion

The zebrafish *atoh7* mutant fails to produce RGCs and therefore provides a unique model in which the brain is entirely disconnected from the retina throughout development and during adulthood.

Although the phenotype of larval *atoh7* mutants has been thoroughly described, data on adult mutants remains sparse ^14,19^. This is likely due to difficulties in raising *atoh7* mutants, which do not survive under standard feeding conditions.

The present study aims to address this shortage by employing an adjusted raising strategy; i.e. providing an excess supply of food, ensuring adequate adult survival rates (Fig. 1A,B). Adult *atoh7* mutants are viable and fertile, while still exhibiting the characteristic phenotype of dark body pigmentation due to complete blindness. However, compared to wildtype and heterozygous siblings, adult *atoh7* mutants are significantly smaller (Fig. 1C,D), likely caused by early development malnutrition which could not be compensated for with excess amounts of food.

### Adult *atoh7* mutants largely retain larval phenotype

The retinal phenotype we observed in adult mutants closely parallels the features documented in larval stages. Kay and colleagues have shown that up until 6 days post-fertilization (dpf), RGC generation is absent in mutants, precluding the formation of an optic nerve ^19^. Instead, the GCL was depopulated of RGCs, with remaining cells found to be misplaced amacrine cells. Likewise in the adult mutant, no RGCs could be identified and consequently neither nerve fiber layer nor optic nerve have been formed (Fig. 2,3). Moreover, we found the adult mutant retina to contain an excess of inner retinal neurons, bipolar and amacrine cells, as well as Müller glia (Fig. 4). Similarly, Kay et al. noted an elevated count of bipolar cells and Müller glia; this increase of cell numbers in larval mutants mirrors also the proportions we observed in adult *atoh7* mutants (2-fold for bipolar cells and a 10-20% increase for Müller glia) ^19^. Notably, we also observed aberrations in the photoreceptors of the adult mutants, with shortened rod outer segments and a disrupted cone mosaic, that have not been described before. Similar to the phenotype described in fish, Brown and colleagues have reported a lack of RGCs and optic nerve as well as decreases in INL and ONL thickness in a math5 (ortholog of *atoh7*) mouse model ^22^. Interestingly, they also observed a disrupted layering of the retina as well as a significant increase in the number of cones. They propose a binary change from RGCs to cones in development which might also occur in fish.

Birth-dating combined with cell marker studies underscored the pivotal role of *atoh7* in RGC determination during the initial phase of retinal neurogenesis, aligning with findings from murine models ^33,34^. Recently it was posited that *atoh7* is not a direct essential competence factor, but rather regulates retinal ganglion axon targeting and survival ^35^. Failure of *atoh7* regulation in turn results in absence of target-derived trophic factors of RGCs without axons in the ON, and subsequent cell death of RGCs ^36,37^. Lineage tracing experiments in larvae revealed that progenitor cells remained undifferentiated for an extended period, with a delay of approximately two cell cycles, bypassing the first wave of neurogenesis ^38^. Consequently, retinal progenitors accumulate, yielding increased numbers of amacrine and bipolar cells during the second wave of neurogenesis. This increase in non-RGC neurons was predicted by a stochastic lineage progression model, resulting in excess total numbers of retinal cells in larval *atoh7* mutants compared to wildtype ^38^.

Additionally, the absent replenishment of RGCs in *lakritz* mutants indicates that *atoh7* is necessary for RGC differentiation. This strengthens the findings of Pittman et al., who transiently decreased RGCs through translation-blocking *atoh7*-morpholinos ^39^. The permanently RGC depleted retinae of the *atoh7* mutant lends itself for further investigation of the development and functional role of RGCs, for example through transplantation experiments of healthy or mutant RGCs ^40^ or to study downstream genes of *atoh7*, like barhl1a ^41^.

### Quantitative OKR assay shows that adult *atoh7* mutants are blind

Larval *atoh7* mutants lack RGCs, and therefore the functional connection between retina and visual processing areas in the brain, and thus are blind ^14,19^. This has been shown via the absence of the OKR, i.e. the failure of larvae to exhibit the stereotypical tracking behavior of moving stimuli. We show that this inability is retained in adult mutants, using a similar, quantitative OKR-based vision test (Fig. 3G) ^31^. Curiously, some of the fish still showed the stereotypical tracking behavior (slow tracking phases interrupted by fast saccades); this ‘residual OKR’ however was not purposeful or directed, but rather occurred independently of the moving stimulus. Wu and colleagues investigated the underlying neuronal circuitry of the OKR in larval zebrafish, and found a cluster of cells in the pretectal M1 area to be required to drive OKR behavior. Moreover, optogenetically activating those cells was sufficient to evoke the OKR in the absence of a visual stimulus ^42^. This pretectal M1 area later develops into the superficial pretectal region in adult zebrafish ^43^ and is involved in higher order visual and multi-sensory processing ^44^. While Wu et al. showed that inhibiting axonal RGC signals to the brain abolishes the OKR in larvae ^42^, we tentatively suggest that in adults, M1 cells are activated also by non-visual input (such as from the functioning vestibular system), which in turn may drive the aimless OKR in *atoh7* mutants.

### Missing retinal input results in structural changes in the optic tectum

The absence of RGCs in the *atoh7* fish, which normally form the retinofugal axons innervating the brain, resulted in no discernible axonal connection between the retina and brain. In wildtype, the vast majority (97%) of RGC axons terminate in the optic tectum ^45^.

Predominantly involved in visual processing, the optic tectum is a complex segment of the mesencephalon ^46,47^, composed of 12 distinct layers ^44,48^. Our measurements of different areas of explanted brains revealed a significant size reduction in the mutant optic tectum (Fig. 5A). Histological analyses further confirmed this size reduction was exclusive to the optic tectum, with the underlying tegmentum remaining unchanged (Fig. 5C). The decrease in mutant optic tectum area was solely attributed to a reduction in white matter, as the area of the periventricular grey zone (PGZ) remained the same size compared to wild type siblings (Fig. 5E). In fact, the PGZ of *atoh7* mutants appeared thicker compared to wildtype, likely a secondary effect of the smaller optic tectum, as the same amount of cell bodies had to be distributed on less lateral space. The tectal ventricles are similarly affected in *atoh7* fish, and due to the smaller optic tectum, ventricle space appears compressed (Fig. 5F).

In wildtype, the stratum opticum is a layer of the optic tectum consisting of the afferent nerve fiber bundles of the ON terminating in the optic tectum ^48^. Expectedly, in mutants no stratum opticum could be observed (Fig. 5G). Moreover, the stratum fibrosum et griseum superficiale (SFGS), recognized as the optic tectum’s primary visual layer ^49^, exhibited significant reduction of thickness. These findings support the notion that lack of retinal innervation results in structural changes in the optic tectum. This is also supported by previous research, indicating that the optic tectum’s size is directly proportional to the amount of retinal input it receives ^50–52^. A putative mechanism for the size reduction could be via missing stimuli-dependent BDNF/TrkB signaling. It was shown that depriving the retina of light for a prolonged time, or the optic tectum of retinal input via optic nerve crush, leads to a reduction of BDNF in the respective tissue ^53,54^ and a reduction of neural stem cells in the optic tectum (Sato et al. 2016). This phenotype could be rescued using a TrkB agonist, suggesting that in the optic tectum NSC proliferation is stimuli dependent ^55^. Shh and Wnt signaling could also be putative contributors to retinal-input dependent NSC proliferation, as it was shown that Shh and Wnt regulate NSC proliferation in the optic tectum in zebrafish ^56^, which could stem from a source transmitting Shh via retinal axons, as in the developing Drosophila brain ^57^. Although our focus remained on the optic tectum, preliminary observations of the 12 retinorecipient nuclei not located in the optic tectum, found no obvious changes. However, we could observe a significant increase in size of the mutant telencephalon (Figure 5B), suggesting, for example, a potential compensation for visual loss through an enhanced olfactory system ^44,58,59^.

### *Atoh7* mutations produce variable phenotypes in humans

For humans, the OMIM database (Online Mendelian Inheritance in Man) currently lists 4 families with mutations in the *atoh7* locus affecting both the coding sequence but also the region upstream of the transcription start site. All affected patients show a similar phenotype that include congenital blindness, nystagmus, glaucoma, corneal opacities, microphthalamus and persistent hyperplastic primary vitreous, caused by a failure of the embryonic vitreous and hyaloid vasculature to regress during development ^23,26,60,61^. A recent study compared the transcriptome of *atoh7* mutant larvae to a wildtype transcriptome to identify potential genetic networks underlying human eye disease ^62^. They identified clusters enriched in retinal development, cell cycle, chromatin remodeling, stress response and Wnt pathway to be differentially expressed in mutants. However, as of now, no follow-up studies on the underlying molecular mechanism that results in these global eye defects as observed in human patients have been performed. This discrepancy between the described phenotype in human and zebrafish mutants can potentially be ascribed to the different underlying mutations. The generation of mutants exactly modelling the human mutations will be of pivotal help for answering these questions.

### The adult *atoh7* mutant provides an ideal platform for further experiments

As a fully blind adult organism that permanently lacks any retinal input to higher visual brain areas, the *atoh7* mutant provides a unique tool for future research with many potential applications. While both *atoh7* and bumper, the other blind adult zebrafish mutant available, can be usefully employed as e.g. negative controls for visual behavior experiments or studying neuronal circuitry systems without any visual input, *atoh7* mutants may also be well suited for developing potential therapeutic approaches for retinal disease, such as NCRNA or glaucoma ^26^.

Glaucoma is a widespread neurodegenerative disease, characterized by the loss of RGCs, resulting in partial or complete blindness ^63,64^, for which there are currently no restorative treatment strategies ^65^. *Atoh7* mutants provide a unique platform to investigate how targeted regeneration ^66^ or transplantation ^40^ of RGCs could mitigate these disease phenotypes. Such investigations might explore, for instance, if and how RGC axons could reach retinofugal targets out of temporal and developmental context, or how rewiring of retinofugal axons in the absence of the ON ^67^ takes place, or how survival of RGCs without target-derived trophic factors ^35^ in the adult fish functions in detail. In summary, the availability of adult fully blind and well characterized *atoh7* mutant zebrafish can provide a versatile model for vision research and the development of therapeutic solutions.

## 5 Materials and methods

### 5.1 Ethical statement

All relevant European Union regulations, German laws (Tierschutzgesetz), the Association for Research in Vision and Ophthalmology (ARVO) statement on the Use of Animals in Ophthalmic and Vision Research, and the NIH *Guide for the Care and Use of Laboratory Animals* (National Academies Press, 2011) were strictly followed for all animal work. Protocols were approved by the Institutional Animal Welfare Officer (Tierschutzbeauftragte(r)) and licensed by the regional Ethical Commission for Animal Experimentation (Landesdirektion Sachsen, Germany; license no. DD24-5131/346/11, DD24-5131/346/12, TVV 59/2018, AV A 2/2022, TVV 21/2018, TV 1/2019). All efforts have been made to minimize animal suffering and number of animals used.

### 5.2 Zebrafish husbandry

Fish were kept under standard conditions and maintained at 26 °C on a 14 h light, 10 h dark cycle as previously described ^29^. We used adult (6 to 12 months of age) *lak^th241^* ^17^ fish as well as their wildtype siblings (herein referred to as *atoh7^+/+^* and *atoh7^-/-^*, respectively). Heterozygous fish displayed a similar phenotype as the *atoh7^+/+^* and are therefore not further mentioned. Both female and male fish were used in equal ratios in all experiments. Body length was measured from the tip of the mouth to the base of the caudal fin. Behavioral experiments were performed during the light cycle at the same time of the day.

### 5.3 Raising of *atoh7* mutants

Heterozygous *lak^th241^* fish were crossed and the resulting offspring was incubated in E3 medium ^29^ at 28 °C. At 6 days post fertilization (dpf), larvae were sorted according to their pigmentation (dark or light) after illumination with a lamp for 2 hours (as described in ^19^). Larvae with dark or light phenotype, respectively, were raised separately albeit under comparable conditions (see Fig. 1A). Until 8 wpf, animals were kept in 200 ml beakers with manual water exchange every other day and exclusively fed with paramecia. Excess amounts of food were critical for the survival of the mutants throughout the first 2 months. At 8 wpf, animals were transferred into mouse cages and additionally fed with artemia and water quality was carefully monitored. At 12 wpf, mouse cages were connected to the recirculating system with automatic water exchange. Feeding of paramecia was terminated and successively replaced with dry food, in addition to paramecia. By 16 wpf, animals were exclusively fed with dry food and moved to tanks.

### 5.4 Genotyping of *atoh7* mutants

A PCR strategy was used to genotype *atoh7* fish (modified from ^19^). In brief, genomic DNA was isolated from fin clips and stored at 8 °C. PCR was performed using primers that amplified a region in the exon of the *atoh7* gene flanking the mutation site (primers: for: 5’- CCG GAA TTA CAT CCC AAG AAC-3’, rev: 5’-GTG TAT GAT ATT CAG CTC TCG ACT-3’, DreamTaq (Thermo Scientific), 53 °C annealing)) (Fig. S1B). The generated PCR product (total length 777 bp) was digested using EcoR147I (Fast Digest, Thermo Scientific). Wildtype fish were characterized by two distinct bands (593 bp and 184 bp), heterozygous *atoh7* fish by three bands (777 bp, 593 bp and 184 bp) and homozygous *atoh7* fish by one band (777 bp) (Fig. S1C). To confirm the genotyping strategy, bands were cut from the gel with a scalpel and DNA was purified using the Nucleospin™ Gel and PCR Clean-up Kit (Machery-Nagel) and sent for sequencing (Eurofins Genomics) (Fig. S1D).

### 5.5 Optical coherence tomography

Optical coherence tomography (OCT) measurements were performed as described in ^31^. Here, a custom-built spectrometer-based OCT setup was used for non-invasive *in vivo* imaging of the adult zebrafish retina enabling an axial and lateral resolution of 1.3 and 7 µm in air, respectively. OCT recordings were performed as described previously ^31,68^.

### 5.6 Optokinetic response test

Measurements of the optokinetic response (OKR) were performed as described previously^31^. Here, a modified commercial system (VisioTracker, TSE Systems GmbH) was employed to generate the stimulus and to measure the optokinetic response (OKR) of adult zebrafish as previously described ^69,70^. Prior to the recordings, fish were presented a training-stimulus (100%, black-white, 0.2 cpd, 15 dps) for up to 5 min to ensure that effects of the anesthesia had subsided and reliable tracking results could be achieved. Then, fish were presented the experimental stimuli (30 % contrast, 15 dps, 0.2/0.3/0.4 cpd). Analysis was conducted with custom made R scripts based on processed data to calculate the average gain (tracking phase velocity to stimulus velocity) for each tested condition.

### 5.7 Histology and immunohistochemistry

HE stainings were performed as described previously ^31^.

For retinal sections, fish were sacrificed using 0.2 % MS-222. Afterwards the corneas were opened and the lenses removed before the skulls were opened carefully. Entire heads were fixed in 4 % PFA overnight at 4 °C and subsequently incubated in EDTA/Sucrose buffer at 4 °C for 24 h. Finally, heads were embedded in 7.5 % Gelatine/20 % Sucrose in 0.1 M Phosphate buffer and stored at - 80 °C. Heads were cut into 14 µm sections (series of 4) using a HM 650 Kryostat (Microm) and collected on glass slides (SuperFrost Plus, Thermo Fisher Scientific). Slides were stored at - 20°C until further usage.

Prior to immunohistochemical stainings, sections were dried at 55 °C for 2 h and washed 1 x 10 min in PBS and 2 x 10 min with PBSTx at RT. For HuC/D stainings an antigen retrieval was performed by incubating the slides for 5 min in 50 mM Tris HCl pH 8.0 at 99 °C before cooling them down to RT and washing 2 x 10 min in PBSTx. Primary antibodies (see table S1) were diluted in PBSTx and incubated overnight in a humidified chamber at 4 °C. Afterwards, sections were washed for 3 x 10 min in PBSTx before incubation with the secondary antibody (see table S2) diluted in PBSTx for 2 h in the dark at RT in a humidified staining chamber. Sections were washed for 10 min with PBSTx and incubated with DAPI (Thermo Scientific, 1:1000 - 5000) in PBSTx for 3 min at RT to counterstain cell nuclei. After final washing in PBSTx, slides were mounted with 70 % Glycerol and stored at 4 °C.

### 5.8 Image acquisition and processing

Imaging was performed at the CMCB Light Microscopy Facility. Microscopic images were acquired with the ZEISS Axio Imager.Z1 microscope equipped with an ApoTome.2 using a Plan-Apochromat with 10x/0.45, 20x/0.8 or 40x/0.95 objectives. An Axiocam MRm (1388×1040, 6.45×6.45 µm, b/w) or an Axiocam MRc (1388×1040 pixel, 6.45×6.45 µm, colour) was used for detection. Sequential image acquisition was used in samples co-stained with multiple fluorophores. For imaging, *z*-stacks were acquired to cover the full information in all three dimensions (typically 10-15 *z*-planes with an interval of 0.4-1.5 µm depending on the numerical aperture of the objective). Close-up images were generated from the original overview image. For overview images, a maximum projection was taken from several *z*- planes. Brightfield images of larvae and adult fish were acquired with a stereomicroscope using Olympus cellSens Dimension (version 1.14) imaging software. Images were processed using Fiji (ImageJ, version 1.52n), ZEN blue 2.6 and Adobe Photoshop CS6. Figures were assembled using PANTONE Affinity Designer (version 1.10.4.1198).

### 5.9 Analysis of gene expression

#### 5.9.1 RT-qPCR

Fish were sacrificed using 0.2 % MS-222, retinae were dissected in ice-cold PBS and immediately frozen in liquid nitrogen. A total of 3 retinae were pooled per sample and stored at - 80 °C until further usage. RNA was extracted from lysed, homogenized tissue with the Total RNA Purification Kit (Norgen Biotek) according to manufacturer’s instructions and reprecipitated if necessary. Intact total RNA was reverse transcribed into cDNA using the Transcriptor First Strand cDNA Synthesis Kit (Roche) according to manufacturer’s instructions with random primers. RT-qPCR was performed after addition of SybrGreen (Roche) (*islet2b* for: 5’- TAT GGG GGA TCA TTC CAA AAA GAA -3’/rev: 5’- GTC GTG GAT CTG ACT GCC G -3’; *actb* for: 5’-CGA GCA GGA GAT GGG AAC C-3’/rev: 5’-CAA CGG AAA CGC TCA TTG C-3’). Expression relative to ß-actin as normalization gene was calculated from Ct values according to the efficiency and delta delta Ct method. At least three biological replicates were analyzed, which were each represented by the average normalized relative ratio of three technical replicates.

#### 5.9.2 *In situ* hybridization

Sections for *in situ* hybridizations were prepared as described before ^71,72^. For generation of the *islet2b* probe, *islet2b* cDNA was amplified from total cDNA using specific primers (for: 5’- GAG CGG GAT ACA AGG CTA CC -3’/rev: 5’- CAG CGG AAG CAT TCG ATG TG -3’), purified from the gel using the Nucleospin™ Gel and PCR Clean-up Kit and cloned into the pCRII-topo vector (Invitrogen) according to the manufacturer’s instructions. The plasmid was transformed into competent *E.coli* XLblue2 and purified using the GeneJET Plasmid Miniprep Kit (Thermo Fisher Scientific). The antisense probe was transcribed with SP6 polymerase using a DIG-labeled NTP mix (Roche Diagnostics).

The *in situ* hybridization was performed as described previously ^71,72^.

### 5.10 Quantification and statistical analysis

For cell counts in the retina, 3 consecutive retinal sections rostrally of the optic nerve were considered per sample and normalized to the length of each individual retina. Then the average of all positive cells per mm retinal length was calculated in each experimental group. For measurements in HE stainings, the freehand lines tool in Fiji (ImageJ, version 1.52n) was used. For OCT cross sections, measurements were always taken in the same retinal region (image 400 out of the image stack consisting of 800 cross sections). Length was measured in pixels before being converted into microns (conversion factor 0.526).

For statistical analysis the GraphPad prism software (version 9.0.0) was used to determine *p*-values with either unpaired t-test, one-way ANOVA with Tukey’s multiple comparison test for post hoc analysis or Dunnett’s multiple comparison test for post hoc analysis against control values. Chosen significance levels were \*\*\*\**p* ≤ 0.0001; \*\*\**p* ≤ 0.001; \*\**p* ≤ 0.01; \**p* ≤ 0.05. Values above *p* > 0.05 were not considered significant. Quantifications are shown as scatter plots including mean ± SD.

## Acknowledgements

We would like to thank past and present members of the Brand lab for many lively discussions as well as Daniela Mögel and Sylvio Kunadt for excellent fish care. In addition, we thank Dr. Judith Konantz and Dr. Heiner Grandel for helpful comments on the manuscript and all members of the Clinical Sensoring and Monitoring Group at TU Dresden, especially Dr. Christian Schnabel and Mirko Mehner for support with the OCT setup. Moreover, we would like to thank Karin Finger-Baier for helpful comments on raising the adult mutants. This work was supported by the Light Microscopy Facility (Dr. Ruth Hans) and Histology Facility (Susanne Weiche), core facilities of the CMCB at the Technische Universität Dresden.

## Figure legends

**Figure S1:**
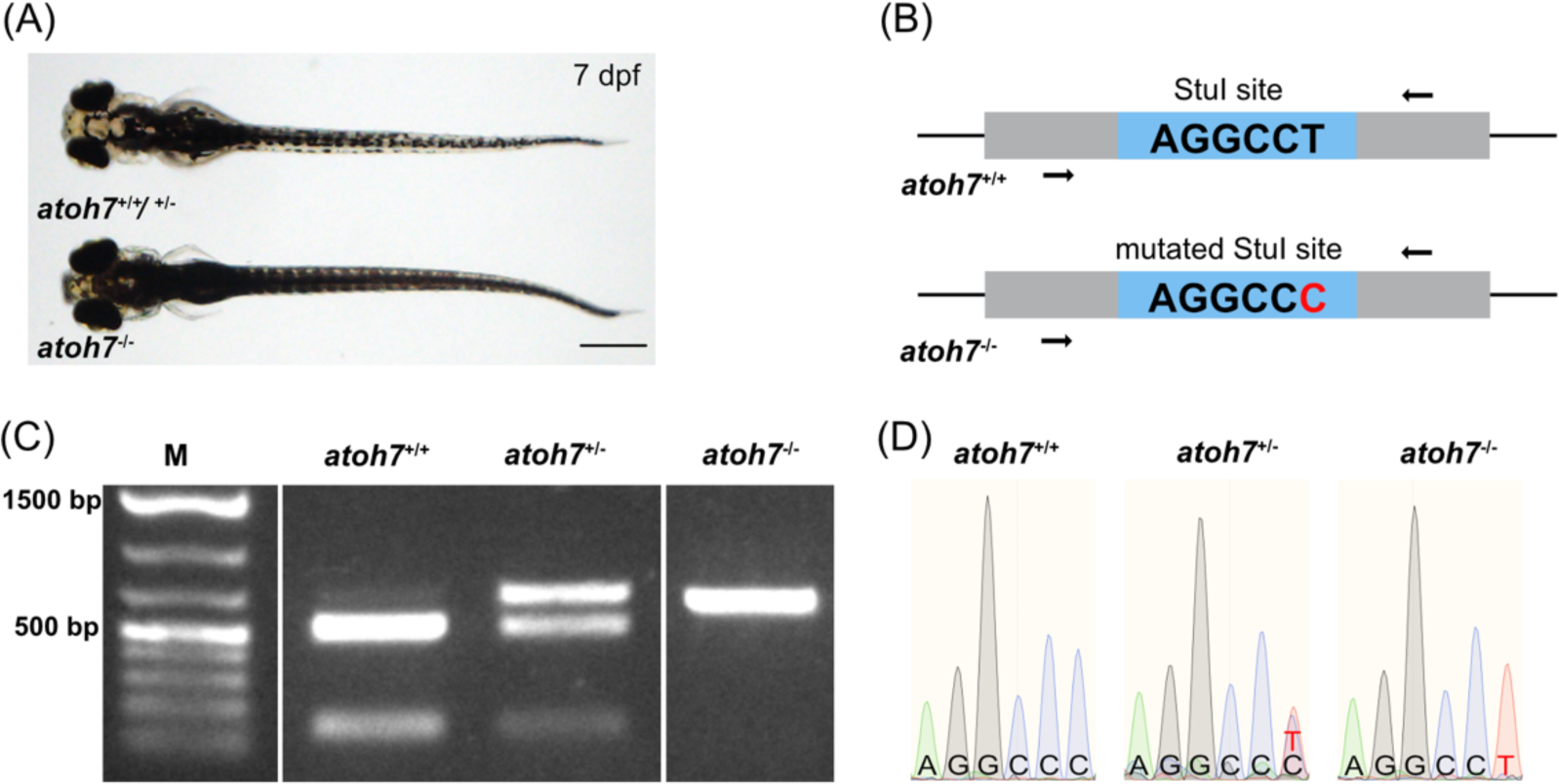
Phenotypic characterization and genotyping of *atoh7* mutants at larval stages. (A) At 7 days post fertilization (dpf), *atoh7*-/- larvae can be distinguished from their wildtype and heterozygous siblings by a darker pigmentation. (B) The underlying mutation is characterized by a single base pair exchange (T to C) resulting in the removal of a StuI restriction site. Arrows indicate primers annealing to up- and downstream genomic regions in wildtype and mutants. (C) Amplification of the genomic regions followed by StuI digestion of the PCR product results in genotype-specific patterns: *atoh7*+/+ 593 + 184 bp; *atoh7*+/- 777 + 593 + 184 bp; *atoh7*-/- 777 bp. M: marker (1 kb plus ladder, Thermo Fisher Scientific)). (D) Sequencing of the respective PCR products confirmed the PCR results.

**Figure S2:**
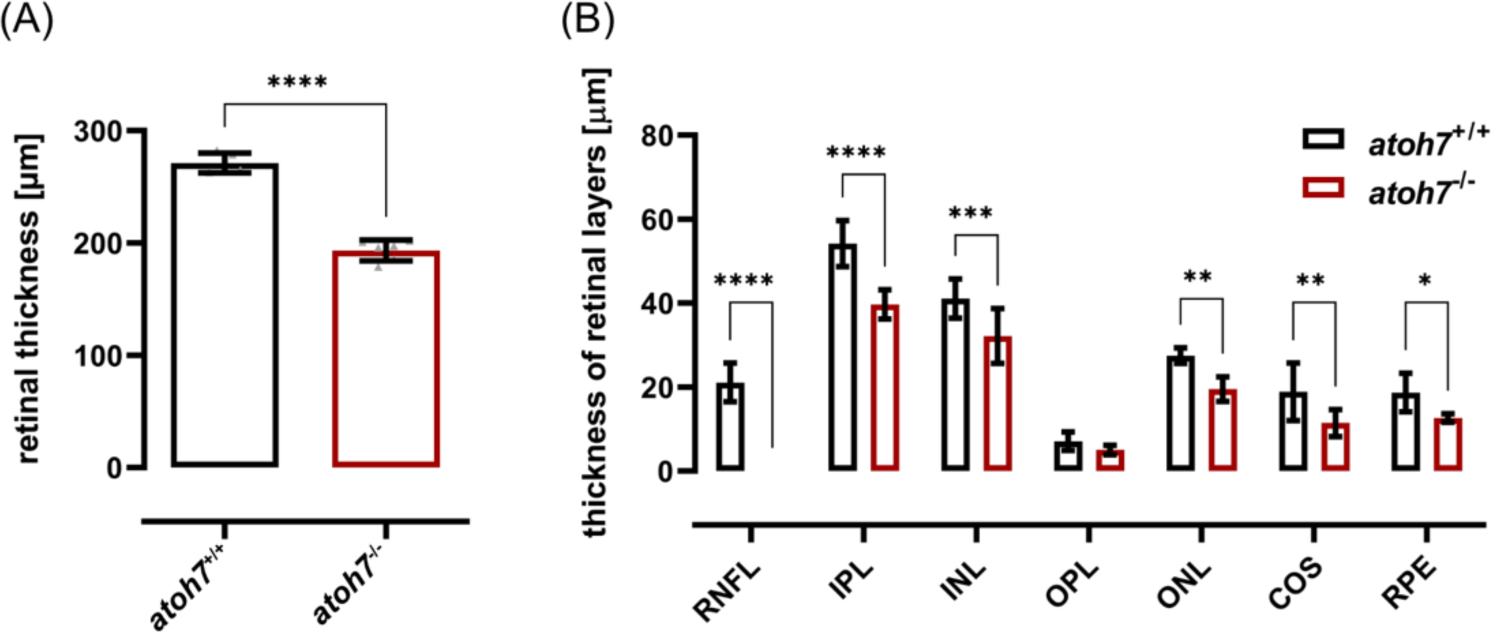
Absolute measurements of whole retinal thickness and individual layers. Quantifications of the absolute thicknesses of the retina (A) and retinal layers (B) measured in in vivo OCT cross sections revealed a significant reduction of the total retinal thickness and all retinal layers except OPL in mutants. The RNFL was not visible and hence could not be measured in the mutants. Statistics: all data are represented as mean ± SD, unpaired t test (B, D), t test with multiple comparisons (C, E), p < 0.05 (*); 0.01 (**); 0.001 (***) or 0.0001 (****). Abbreviations: COS – cone outer segments, GCL – ganglion cell layer; INL – inner nuclear layer, IPL – inner plexiform layer; ONL – outer nuclear layer, OPL – outer plexiform layer; RNFL – retinal nerve fiber layer; ROS – rod outer segments, RPE – retinal pigment epithelium.

**Figure S3:**
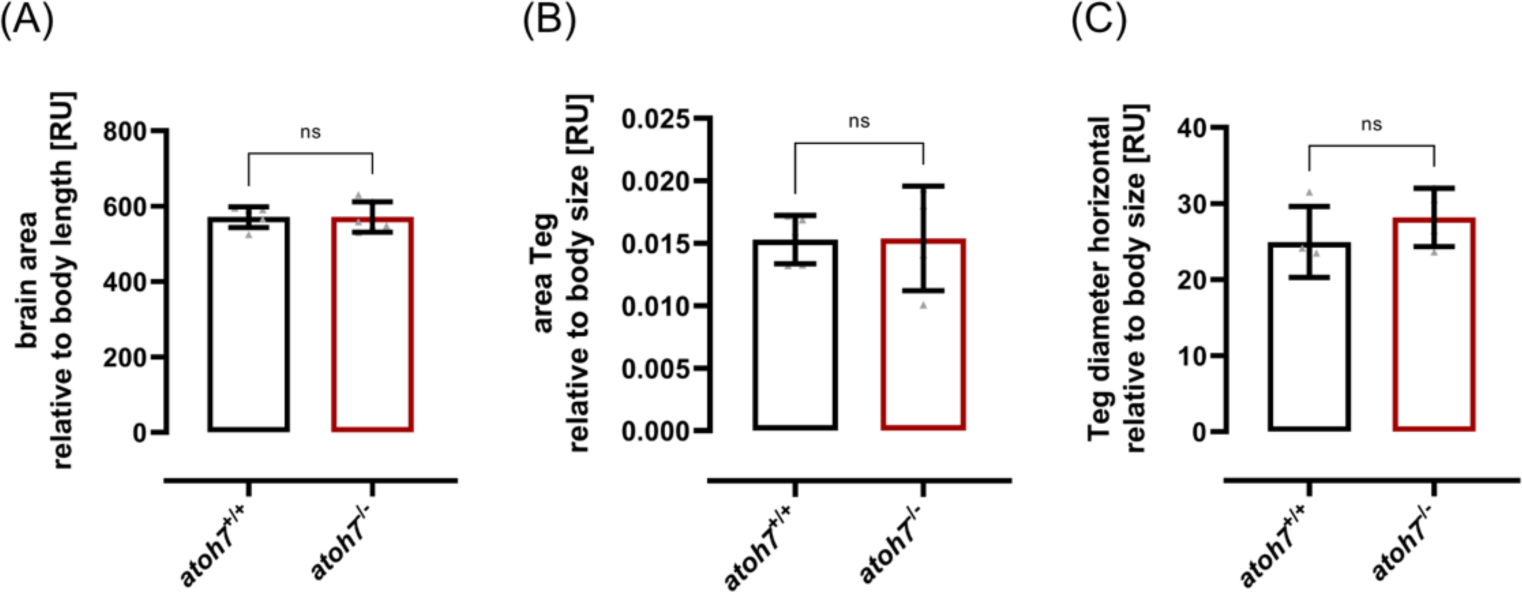
Quantifications of brain areas. (A) Quantifications of the whole brain area relative to body size resulted in no statistical difference between wildtype and mutants. (B,C) Area of the tegmentum (Teg) normalized to body size (B) as well as tegmentum diameter normalized to body size (C) is not statistically significant between wildtype and mutants. Statistics: all data are represented as mean ± SD, unpaired t test, p < 0.05 (*); 0.01 (**); 0.001 (***) or 0.0001 (****).

## Tables

**Table S1:**
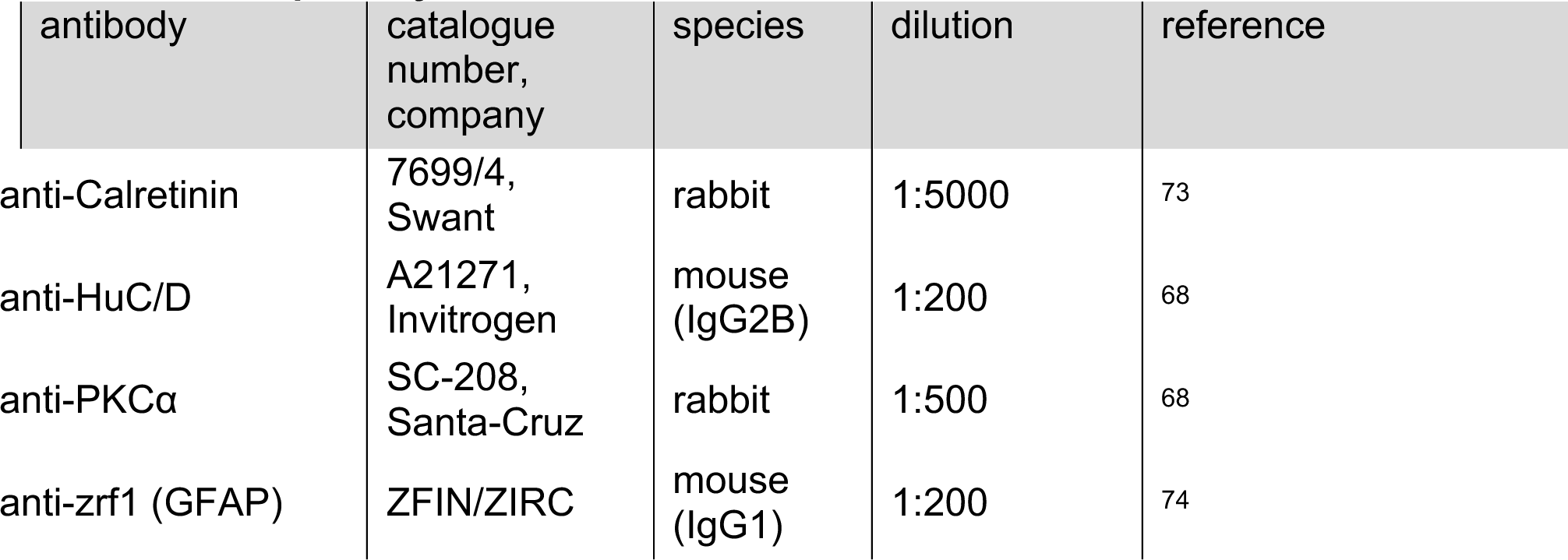
List of primary antibodies.

| antibody | catalogue number, company | species | dilution | reference |
| --- | --- | --- | --- | --- |
| anti-Calretinin | 7699/4, Swant | rabbit | 1:5000 | 73 |
| anti-HuC/D | A21271, Invitrogen | mouse (IgG2B) | 1:200 | 68 |
| anti-PKCα | SC-208, Santa-Cruz | rabbit | 1:500 | 68 |
| anti-zrf1 (GFAP) | ZFIN/ZIRC | mouse (IgG1) | 1:200 | 74 |

**Table S2:**
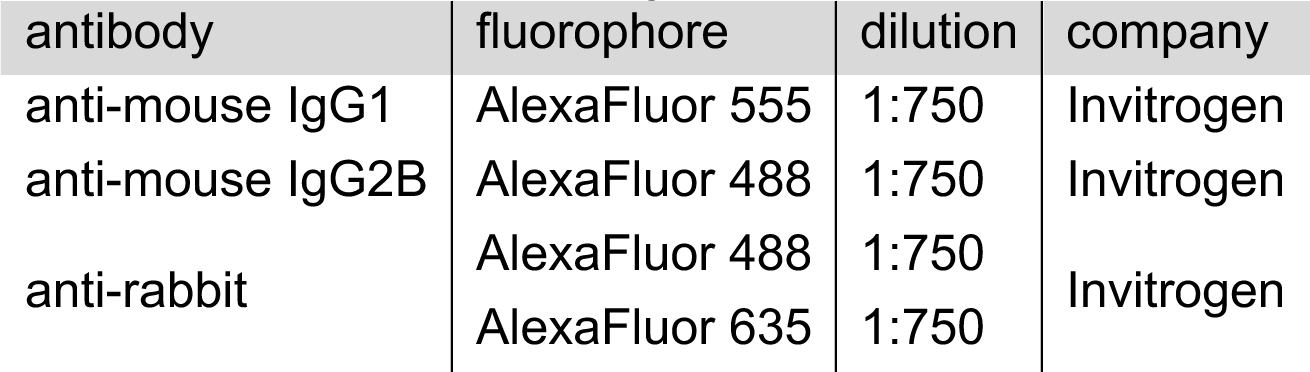
List of secondary antibodies.

**Table S3:**
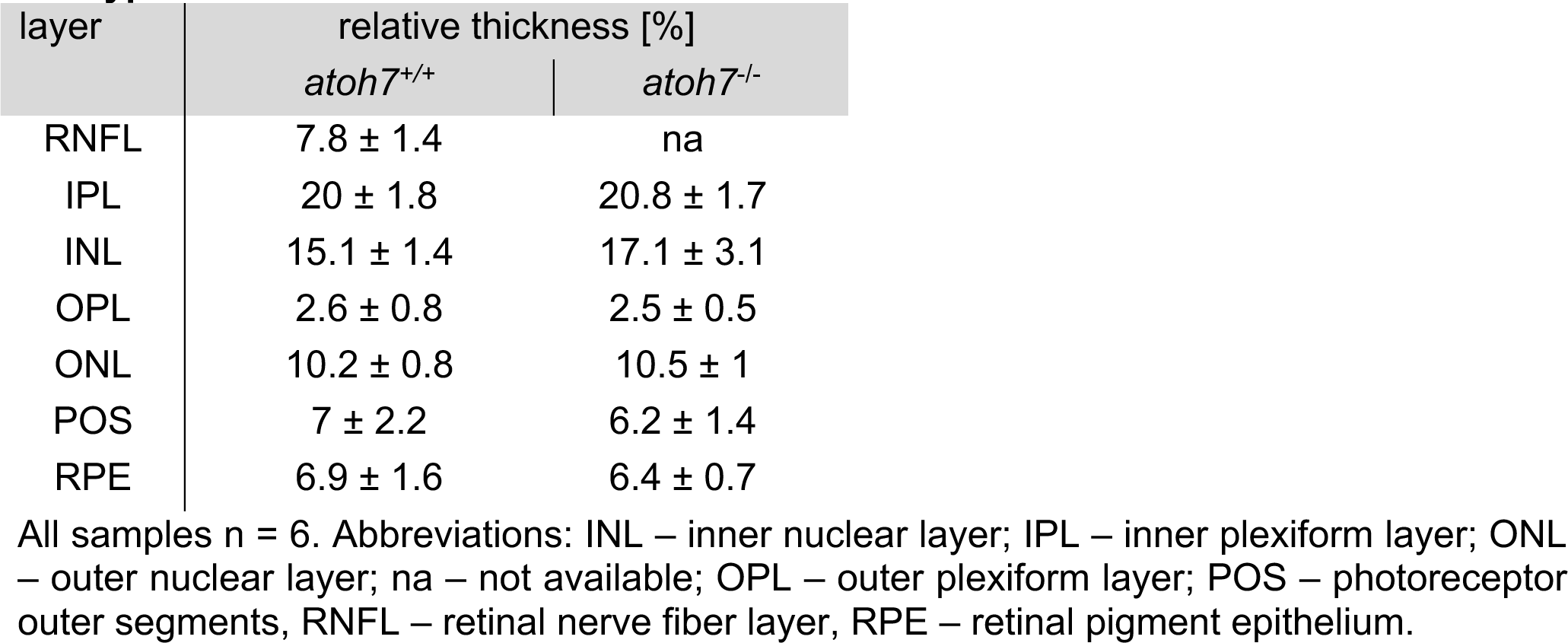
Absolute and relative thicknesses of retinal layers in *atoh7* mutants and wildtypes.

| layer | relative thickness [%] |  |
| --- | --- | --- |
|  | *atoh7*<sup>+/+</sup> | *atoh7*<sup>-/-</sup> |
| RNFL | 7.8 ± 1.4 | na |
| IPL | 20 ± 1.8 | 20.8 ± 1.7 |
| INL | 15.1 ± 1.4 | 17.1 ± 3.1 |
| OPL | 2.6 ± 0.8 | 2.5 ± 0.5 |
| ONL | 10.2 ± 0.8 | 10.5 ± 1 |
| POS | 7 ± 2.2 | 6.2 ± 1.4 |
| RPE | 6.9 ± 1.6 | 6.4 ± 0.7 |

All samples n = 6. Abbreviations: INL – inner nuclear layer; IPL – inner plexiform layer; ONL – outer nuclear layer; na – not available; OPL – outer plexiform layer; POS – photoreceptor outer segments, RNFL – retinal nerve fiber layer, RPE – retinal pigment epithelium.

